# *Wolbachia* endosymbionts in two *Anopheles* species indicates independent acquisitions and lack of prophage elements

**DOI:** 10.1101/2021.11.15.468614

**Authors:** Shannon Quek, Louise Cerdeira, Claire L Jeffries, Sean Tomlinson, Thomas Walker, Grant L. Hughes, Eva Heinz

## Abstract

*Wolbachia* is a genus of obligate bacterial endosymbionts that infect a diverse range of arthropod species as well as filarial nematodes, with its single described species, *Wolbachia pipientis,* divided into several ‘supergroups’ based on multilocus sequence typing. *Wolbachia* strains in mosquitoes have been shown to inhibit the transmission of human pathogens including *Plasmodium* malaria parasites and arboviruses. Despite their large host range, *Wolbachia* strains within the major malaria vectors of the *Anopheles (A.) gambiae* and *A. funestus* complexes appear at low density based solely on PCR-based methods. Questions have been raised as to whether this represents a true endosymbiotic relationship. However, recent definitive evidence for two distinct, high-density strains of supergroup B *Wolbachia* within *A. demeilloni* and *A. moucheti* has opened exciting possibilities to explore naturally occurring *Wolbachia* endosymbionts in *Anopheles* for biocontrol strategies to block *Plasmodium* transmission. Here we utilise genomic analyses to demonstrate that both *Wolbachia* strains have retained all key metabolic and transport pathways despite their smaller genome size. We further confirm the presence of cytoplasmic incompatibility factor genes, despite noticeably few prophage regions. Additionally, phylogenetic analysis indicates that these *Wolbachia* strains may have been introduced into these two *Anopheles* species via horizontal transmission events, and unlikely to be by ancestral acquisition and subsequent loss events in the *Anopheles gambiae* species complex. These are the first *Wolbachia* genomes that enable us to study the relationship between natural strains *Plasmodium* malaria parasites and their *Anopheline* hosts.

**Impact statement:** *Wolbachia* naturally infects a wide range of arthropod species, including insect vectors of human pathogens, where they may play a role in inhibiting their replication. These bacteria have been commonly found within *Aedes (Ae.) albopictus* and *Culex pipiens* mosquitoes but have been noticeably absent in the *Anopheles* mosquito genera, which includes all species responsible for malaria transmission. Recent PCR-based methods have suggested the potential for natural *Wolbachia* strains within the *A. gambiae* species complex, which includes major malaria vector species including *A. gambiae* s.s., *A. coluzzii* and *A. arabiensis*. We recently reported the presence of stable *Wolbachia* strains naturally occurring within two different *Anopheles* species (*A. demeilloni* and *A. moucheti)*. In this study, we perform comparative genomic analysis of these two *Wolbachia* genomes against each other and published *Wolbachia* strains. The current assemblies are some of the smallest sequenced *Wolbachia* strains of insects, although their metabolic pathway repertoire is comparable to other strains. Interestingly, prophage fragments were identified within only one of the two strains. The findings of this study will be of significant interest to researchers investigating *Wolbachia* as a potential malaria biocontrol strategy, giving greater insight into the evolution and diversity of this obligate intracellular endosymbiont.

**Data summary:** Sequence data generated and used for this analysis are available in the National Centre for Biotechnology Information Sequence Read Archive (NCBI SRA bioproject number PRJNA642000). The two assembled *Wolbachia* genomes are available with genome accession numbers GCA_018491735.2 and GCA_018491625.2. Additional *Wolbachia* genomes used for comparative analysis are described in the supplementary material.

The authors confirm all supporting data, code and protocols have been provided within the article or through supplementary data files. Additional supplementary data files used to generate several figures can be found at: https://figshare.com/projects/Wolbachia_endosymbionts_in_two_Anopheles_species_indicates_independent_acquisitions_and_lack_of_prophage_elements/126533

## Introduction

*Wolbachia* has a wide host range, including insects [1] where various estimates have predicted between 52% to 60% of all arthropod species naturally infected [2, 3]. Attempts to characterise the within-species diversity has resulted in the designation of *Wolbachia* ‘supergroups’ A through to T [4, 5], with several exceptions [6], via multilocus sequence typing (MLST) of five single-copy conserved genes [7]. The relationship between *Wolbachia* and their hosts can range from obligate mutualism, where the endosymbiont is essential for host survival and reproduction [8, 9], to reproductive parasitism, where it manipulates the reproduction of its host to spread through the population. Currently, the best-studied phenotype (which also affects mosquito hosts) is cytoplasmic incompatibility (CI), which causes infected males to produce unviable offspring unless they mate with an infected female, while infected females have viable offspring regardless of the males infection status, thus conferring a fitness advantage to *Wolbachia*-infected females.

Genetic studies have previously identified a pair of cytoplasmic incompatibility factor genes, *cifA* and *cifB* [10, 11], that have been correlated to this phenotype. These genes have often been found to co-occur as a single operon within prophage Eukaryotic Association Modules (EAMs), and are believed to spread via horizontal transmission between *Wolbachia* strains due to their localisation within prophage regions [12]. Despite being part of the same operon, these genes have been observed to be differentially regulated, with *cif*A having higher expression relative to *cif*B [13]. When these CI genes were first identified, they were placed into three distinct phylogenetic groups. While all three were recognised to maintain protein domains that had predicted nuclease activity, the catalytic residues for these nuclease domains were predicted to be absent in one of the three groups [10], which instead contained an additional protein domain with ubiquitin-like specific protease activity [11, 14, 15]. This was later characterised as the Type I group [14]. Additionally, recent research has identified genes encoding similar features in other members of the *Rickettsiales* order, often found associated with mobile genetic elements, such as plasmids [14, 16]. As a result, a recent study has identified up to five phylogenetic types, with one of these types being identifiable in other *Rickettsia* as well as *Wolbachia* [14].

Utilisation of the CI phenotype has been explored as the basis for potential mosquito control strategies to reduce human disease transmission. The bacterium is capable of inducing CI in both natural [5, 17, 18] and artificially infected lines [19–21], and possible methods to utilise them for mosquito population control include release of males infected with *Wolbachia* [22], or potentially via release of genetically modified mosquitoes that carry the CI genes, but not *Wolbachia* [23, 24]. In addition to inducing the CI phenotype, *Wolbachia* has been shown to interfere with pathogen replication directly, both in those that cause disease in the insect, as well as human pathogens that utilise the insect as a vector [5, 25–27]. This has been observed to be most effective with artificial infections of *Wolbachia* in non-native host mosquitoes [19–21]. Recent trials have shown that *Wolbachia* can be used to great effect in preventing the spread of dengue virus [28, 29], while laboratory trials have indicated their potential to block *Plasmodium* replication in artificially infected *Anopheles* mosquitoes [30–32]. While there is little evidence to date of stable *Wolbachia* infections [30], a stable infection within *A. stephensi* is possible [31, 32]. Infection of these mosquitoes with *Wolbachia* was associated with significantly reduced hatch rates however [31, 33], possibly affecting the viability of CI as a control tool in this system.

Despite *Wolbachia*’s presence in a wide variety of insects, natural high-density strains within the *Anopheles* genus of mosquitoes have not been conclusively proven [34, 35] until recently [36]. Previous efforts to detect this bacterium required highly sensitive PCR techniques [37–41] that amplify a select handful of *Wolbachia* genes. Unfortunately, this alone cannot confirm the presence of live bacteria or stable *Wolbachia* strains within insects. Furthermore, phylogenetic placement of these amplified *Wolbachia* sequences within *A. gambiae* shows multiple strains distributed across supergroups A and B, with some strains not assigned to any supergroup [35].

We recently demonstrated high-density *Wolbachia* strains in two *Anopheles* species, *A. demeilloni* and *A. moucheti* [36, 42], which we observed in wild populations collected over a large geographic range in temporally-distinct populations. Importantly, we further visualized these bacteria in the germline, as well as sequenced near-complete *Woblachia* genomes from both host species [36]. Here, we present reassembled and scaffolded genomes for both strains, as well as in-depth comparative analyses of these two *Wolbachia* strains against each other, as well as in the broader context of *Wolbachia* supergroups A through to F, with specific focus on supergroup B. We show that, in terms of both size and predicted protein-coding genes, both assembled genomes are at the low end of the range of *Wolbachia* strains found within insects whilst containing reduced or, in the case of *w*AnM, no prophage WO regions. Despite this, both *Wolbachia* genomes maintained complete pathways that are expected for the genus, such as complete haem and nucleotide biosynthetic pathways and type IV secretion systems. Additionally, we reconstructed the phylogenetic history using whole genome sequence data which indicates that these strains may originate from independent acquisitions via horizontal transfer events, and not from an ancestral infection that has since been lost in other *Anopheles* mosquitoes.

## Results

### Assembled *Wolbachia* genomes are small in size but supported by high completeness scores

As *Wolbachia* is an obligate intracellular endosymbiont, they have a highly reduced genome and can only be isolated from infected host material, posing a challenge to obtain complete, uncontaminated genome sequences. The updated genome assembly of *Wolbachia* of *A. demeilloni* (*w*AnD) has a total length of 1,231,247 base pairs (bp), while *Wolbachia* of *A. moucheti* (*w*AnM) has a genome length of 1,121,812 bp. While these genome sizes are smaller compared to other analysed *Wolbachia* strains that reside within insects (particularly *w*AnM), they are larger than the genomes of those found in filarial nematodes, which have a maximum size of 1.08 Mbp amongst those compared in our analysis (Table 1). Refseq annotation of both *w*AnD and *w*AnM genomes identified 1,157 and 1,082 protein-coding genes and 122 and 80 pseudogenes, respectively (Table 1). For comparison, the *Wolbachia* strains of *Ae. albopictus* (*w*AlbB) and *Cx. quinquefasciatus* (*w*Pip) maintained 1,180 and 1,241 protein-coding genes respectively.

**Table 1:** Summary table of a selection of different near-complete *Wolbachia* genomes and their general genome properties. Note the genomes of *w*AnD and *w*AnM (black box highlight), the similar genome properties compared to other *Wolbachia* genomes, but their relatively lower protein-coding gene number when compared against *Wolbachia* strains of supergroups A and B. BUSCO scores were calculated using the Rickettsiales_odb10 lineage, created on 2020-03-06 with a marker gene list total of 364.

| Strain<br>Name | Host organism | Supergroup | Size<br>(Mb) | GC% | Total<br>genes | Proteins | Pseudogenes | tRNA | rRNA | Other<br>RNAs | Ankyrin<br>proteins | BUSCO Score<br>(out of 364) |
| --- | --- | --- | --- | --- | --- | --- | --- | --- | --- | --- | --- | --- |
| **wAu** | *Drosophila simulans* | A | 1.27 | 35.22% | 1,265 | 1,099 | 125 | 34 | 3 | 4 | 35 | 362 (99.5%) |
| **wHa** | *Drosophila simulans* | A | 1.30 | 35.09% | 1,242 | 1,110 | 91 | 34 | 3 | 4 | 36 | 362 (99.5%) |
| **wMel** | *Drosophila melanogaster* | A | 1.27 | 35.23% | 1,247 | 1,144 | 103 | 34 | 3 | 4 | 27 | 361 (99.2%) |
| **wRi** | *Drosophila simulans* | A | 1.45 | 35.16% | 1,340 | 1,245 | 95 | 35 | 3 | 4 | 33 | 360 (98.9%) |
| **wAlbB** | *Aedes albopictus* | B | 1.49 | 34.50% | 1,442 | 1,180 | 221 | 34 | 3 | 4 | 38 | 355 (97.5%) |
| **wAnD** | *Anopheles demeilloni* | B | 1.23 | 33.58% | 1,320 | 1,157 | 122 | 34 | 3 | 4 | 55 | 360 (98.9%) |
| **wAnM** | *Anopheles moucheti* | B | 1.12 | 33.59% | 1,203 | 1,082 | 80 | 34 | 3 | 4 | 37 | 360 (98.9%) |
| **wMa** | *Drosophila mauritiana* | B | 1.27 | 34.00% | 1,196 | 1,055 | 100 | 34 | 3 | 4 | 49 | 360 (98.9%) |
| **wMau** | *Drosophila mauritiana* | B | 1.27 | 34.00% | 1,194 | 1,054 | 99 | 34 | 3 | 4 | 49 | 361 (99.2%) |
| **wMeg** | *Chrysomya megacephala* | B | 1.38 | 33.95% | 1,268 | 1,116 | 111 | 34 | 3 | 4 | 52 | 363 (99.7%) |
| **wNo** | *Drosophila simulans* | B | 1.30 | 34.01% | 1,208 | 1,062 | 105 | 34 | 3 | 4 | 53 | 363 (99.7%) |
| **wPip** | *Culex quinquefasciatus* | B | 1.48 | 34.19% | 1,385 | 1,241 | 103 | 34 | 3 | 4 | 63 | 362 (99.5%) |
| **wOo** | *Onchocerca ochengi* | C | 0.96 | 32.07% | 733 | 645 | 47 | 34 | 3 | 4 | 2 | 346 (95.1%) |
| **wOv** | *Onchocerca volvulus* | C | 0.96 | 32.07% | 734 | 648 | 45 | 34 | 3 | 4 | 3 | 345 (94.8%) |
| **wBm** | *Brugia malayi* | D | 1.08 | 34.18% | 1,029 | 845 | 143 | 34 | 3 | 4 | 18 | 357 (98.1%) |
| **wFol** | *Folsomia candida* | E | 1.80 | 34.35% | 1,662 | 1,541 | 79 | 35 | 3 | 4 | 94 | 362 (99.5%) |
| **wCle** | *Cimex lectularius* | F | 1.25 | 36.25% | 1,238 | 1,023 | 174 | 34 | 3 | 4 | 42 | 356 (97.8%) |

As an obligate intracellular endosymbiont that may have multiple strains infecting the same host, assessing *Wolbachia* genome completeness is important to ensure contaminating reads from different strains are not incorporated, and that the assembly does not have significant gaps. Despite the smaller number of protein-coding genes and genome size, both *w*AnD and *w*AnM were noted to contain over 98% of essential single-copy genes as determined by the BUSCO program [43] (Table 1), with only *w*AnM predicted to have one duplicated gene, indicating that both their respective hosts are infected with only a single strain of *Wolbachia*. These figures are in line with previously published and complete *Wolbachia* genomes of strains found within insects (Table 1), with examples such as the *Wolbachia* strains of *Drosophila* flies (*w*Mel, *w*Ri, *w*Ha, *w*Au and *w*No), and mosquitoes (*w*AlbB, *w*Pip) all having completeness scores ranging from 97.5% to 99.5%. Additional details on these genomes and their associated publications used for comparison are available in Supplementary Table 1.

### Different *Anopheles* species show potentially independent *Wolbachia* acquisition events

Whole-genome phylogenetic analysis was performed to better understand how *w*AnD and *w*AnM may have been acquired by *Anopheles*, utilising the most closely related genome of *Wolbachia* of *Drosophila (D.) simulans* strain Noumea (*w*No) as a reference [36]. Using a total of 36 genomes of *Wolbachia* strains from supergroup B, a single-nucleotide variant (SNV) alignment of 2,824 base-pairs was generated. The midpoint-rooted tree of the SNV alignment (Figure 1) placed both *w*AnD and *w*AnM within a clade that also includes *w*No, and several *Wolbachia* strains that infect *D. mauritiana* [44–46]. We observed a significant number of differences in this alignment between *w*AnD and *w*AnM strains, with a total of 824 SNVs between the two *Anopheles*-derived strains. By contrast, *w*AnM was shown to have a total of 408 and 417 SNVs shared between it and the *Wolbachia* strains of *D. mauritiana* and *w*No respectively, suggesting that *w*AnM is more closely related to these strains than to *w*AnD. Additionally, it was observed that *w*AlbB and *w*Pip, two known *Wolbachia* strains of mosquitoes, do not cluster together, and appear in clades separate from both *w*AnM and *w*AnD. This lack of host clustering can be seen throughout the generated phylogenetic tree (Figure 1), with Insecta host members from different orders appearing throughout. Exceptions to this observation come from *Wolbachia* genomes that have been sequenced from the same host, e.g. *Diaphorina citri* or *Drosophila mauritiana*. Such observations are similar to that in previous studies that predict how *Wolbachia* is not solely restricted to vertical transmission ([47, 48]) and could be an indication of independent horizontal acquisition of *w*AnD and *w*AnM in their current hosts, rather than an ancestral infection that has since been lost in other Anopheline mosquitoes. Phylogenetic analysis of COII and ITS2 sequences of *A. demeilloni* and *A. moucheti* indicate significant phylogenetic distances from both the *A. gambiae* and *funestus* complexes [36]. Furthermore, this study also provided no evidence of resident *Wolbachia* strains within *A. marshallii*, a mosquito species closely related to *A. demeilloni* and *moucheti* [36].

**Figure 1:**
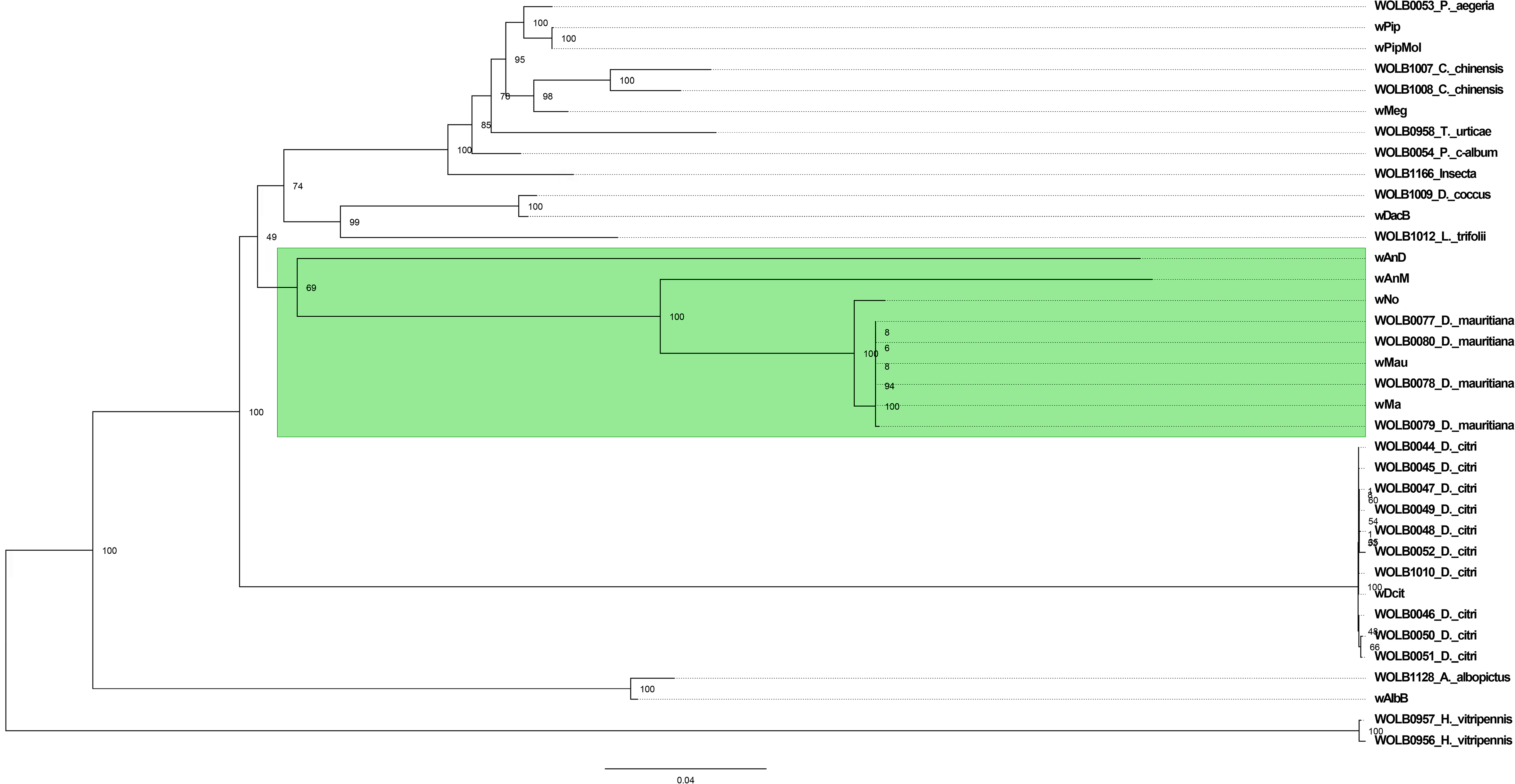
Maximum likelihood phylogenetic tree of whole-genome alignments of a selection of *Wolbachia* genomes, using 1000 bootstrap replicates. Genomes beginning with WOLB followed by four digits were assembled by [44]. Other genomes, with the exception of *w*AnM and *w*AnD (which are bolded and underlined) are the results of previous sequencing efforts, with acronyms as described in Table 1. Tree is midpoint-rooted. Note how *w*AnM and *w*AnD are present within a clade alongside *w*No and several assembled genomes of *Wolbachia* from *Drosophila mauritiana,* (green highlight). By contrast, previously sequenced *Wolbachia* of mosquitoes *w*Pip/*w*PipMol, and *w*AlbB, are present in separate clades.

### The *Wolbachia* core genome is conserved in *w*AnM and *w*AnD orthogroup analysis

Orthologous gene groups are important to identify in *Wolbachia* strains due to their wide distribution across supergroups and diverse hosts, whilst offering insights into the presence/absence of unique pathways that may be involved in host-bacterial symbiosis. For this, we compared the RefSeq annotations of *w*AnD and *w*AnM genomes against 17 *Wolbachia* genomes (Table 1). A total of 18,404 genes were analysed, with 96.8% of these assigned to 1,300 orthogroups, and the remainder left unassigned to any orthogroup. Across the 17 *Wolbachia* strains analysed, a core genome of 9,031 genes distributed across 523 orthogroups was identified (i.e. 40.2% of all identified orthogroups comprising 49.1% of total genes analysed can be considered as part of the core genome, defined as the genes and their protein products that are present in all analysed genomes), with 501 of these orthogroups containing single-copy genes. Outside of this core genome, the number of shared orthogroups is noticeably lower (Figure 2a), and no orthogroups were unique to *Wolbachia* supergroup B strains. For *w*AnD, 18 genes were not assigned to an orthogroup, and no species-specific orthogroups (paralogues present in only one species) were identified (Figure 2a inset). By contrast, *w*AnM was noted to have two species-specific orthogroups containing a total of 47 genes, as well as 13 unassigned genes. None of the protein products for these genes had identifiable protein domains. Two orthogroups containing single-copy genes were identified that was specific to both *w*AnD and *w*AnM, although again none of these had identifiable protein domains.

**Figure 2:**
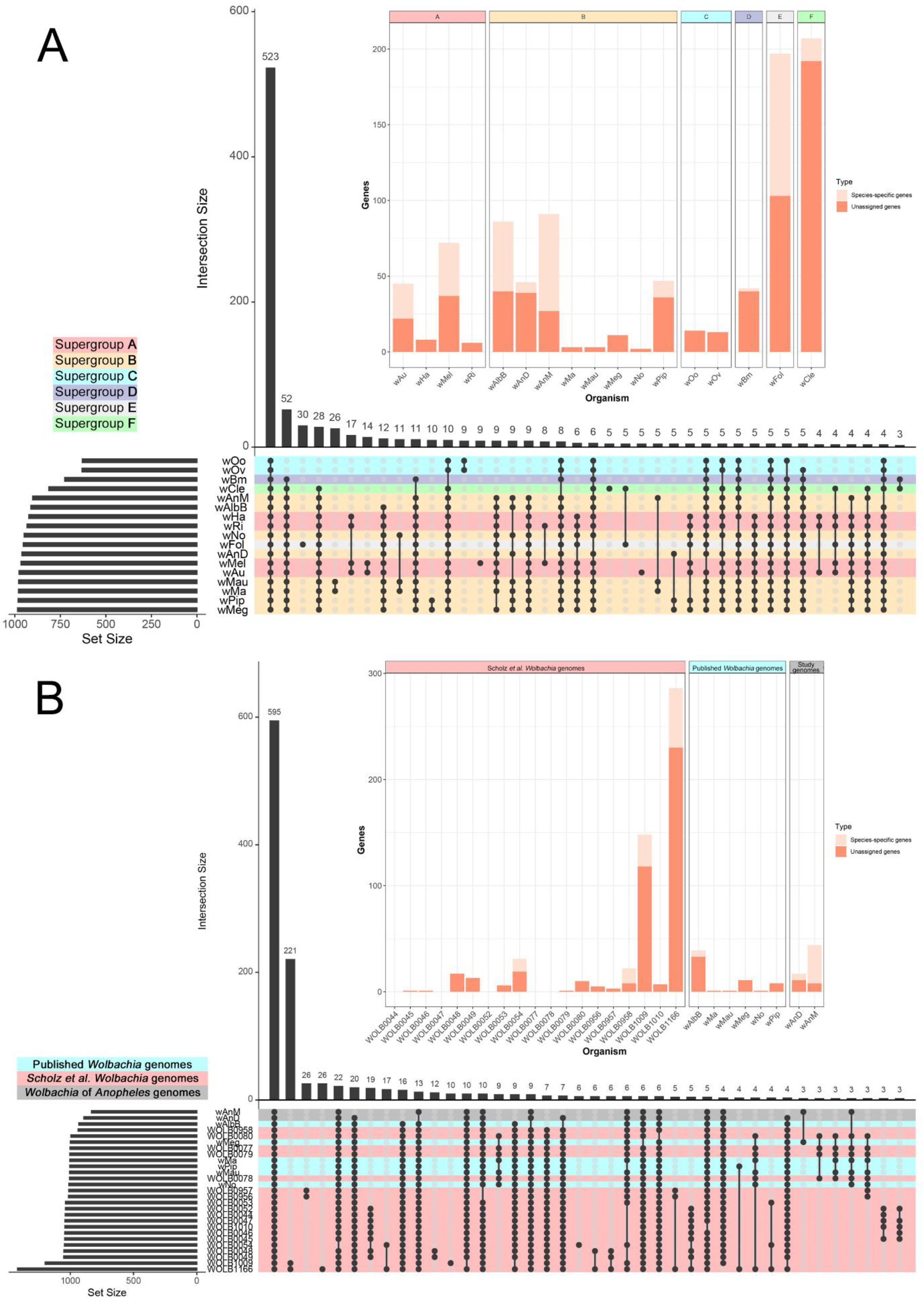
Overview of identified orthogroups amongst *Wolbachia*. **(A)** Graphical representation of set notation of 17 near-complete *Wolbachia* genomes from six of the main supergroups using UpsetR, and the protein orthologues that they encode. Each genome (one per row at the bottom half of the image) is treated as a ‘set’ containing a certain number of orthogroups (denoted by the bar graph on the bottom left of the image). The various permutations of intersections are denoted by the ball-and-stick diagram at the bottom of the image, and the size of these intersections denoted by the bar graph at the top of the image. *Wolbachia* genomes are colour-coded based on their supergroup organisation, with *w*AnM and *w*AnD sets highlighted by a red outline with no fill. Note how an intersect of all 17 *Wolbachia* genomes was identified as containing the vast majority of orthogroups- a ‘core’ proteome total of 523 orthogroups (first bar from left). All other subsequent permutations of intersects contain less than 53 orthogroups. There were no intersects that uniquely contained *w*AnM and *w*AnD, or uniquely contained only supergroup B *Wolbachia*. The inset stacked bar chart shows the distribution of singleton, i.e. genes that do not belong to an orthogroup, dark red bar segment, and strain-specific orthogroups, i.e. genes that belong to a orthogroup unique to that *Wolbachia* strain, light red segment. **(B)** Graphical representation of set notation of 27 supergroup B *Wolbachia* genomes, using the format as described for **(A)**. *Wolbachia* genomes includes eight existing published complete genomes, and 19 recently assembled genomes from Scholz et al., alongside the two recently assembled genomes *w*AnM and *w*AnD (highlighted with a red box). Analysis was performed on local PGAP annotations of all 27 *Wolbachia* genomes. Note how the intersect of all 27 *Wolbachia* genomes shows a core proteome of 595 orthogroups, with the second largest intersect containing 221 orthogroups shared between the Scholz et al. genome assembly for *Wolbachia* of *D. coccus* and an unidentified Insecta. There were no intersects that uniquely contained *w*AnD and *w*AnM.

Further comparisons were performed using a wider selection of *Wolbachia* supergroup B strains, including eight *Wolbachia* genomes used in the previous analysis, as well as a further 19 draft genomes [44] with over 80% completeness. For consistency, all genomes were annotated using a local installation of NCBI’s Prokaryotic Genome Annotation Pipeline (PGAP, [49]). A total of 31,943 genes annotated across the 27 genomes were used, of which 98.4% of these were assigned to 1,678 orthogroups (Figure 2b). A core genome (genes and their protein products that are present in all analysed genomes) was identified containing 15,208 genes distributed across 618 orthogroups (47.6% of total genes were assigned to 36.8% of all orthogroups). Of these 618 orthogroups, 177 contain single-copy genes. A total of 34 and 21 genes were not assigned to an orthogroup for *w*AnD and *w*AnM respectively. One species-specific orthogroup was identified in *w*AnD (containing seven genes), and two species-specific orthogroups were identified in *w*AnM (containing 52 genes). Similar to the previous comparison, one orthogroup was identified as specific to both *w*AnD and *w*AnM, containing single-copy orthologues from both genomes that did not have any identifiable protein domains.

It was interesting to see that the number of orthogroups that could be considered as part of the core genome is less than 50% for both comparisons conducted here. We observed a total of 90 orthogroups that are not considered ‘core’ due to their absence within *Wolbachia* strains of filarial nematodes from supergroups C and/or D specifically, whilst supergroup F strains has 30 unique orthogroups. Additionally, the genomes of *Wolbachia* from *D. mauritiana* (*w*Ma and *w*Mau in Figure 2a) shared 26 unique orthogroups. This observation of an extensive accessory genome has been reported in the past, even among closely-related *Wolbachia* strains [50].

### Despite smaller genomes, the *Wolbachia* spp. core metabolic pathways are conserved in *w*AnD and *w*AnM

Orthogroup analysis of *w*AnD and *w*AnM indicated a high degree of conservation of supergroup B metabolic capacity, and to confirm this, the KEGG Automatic Annotation Server (KAAS, [51]) was used to assign KEGG Orthology. A total of 677 and 660 protein-coding genes were assigned a KO number for *w*AnD and *w*AnM respectively. Subsequent visualisation and manual annotation identified complete biosynthetic pathways that have previously been considered of interest with respect to *Wolbachia*-host symbiosis (Figure 3a). This includes pathways for riboflavin, purines, pyrimidines and haem biosynthesis; and showed all pathways as present in other supergroup B isolates’ genomes. Additionally, both *w*AnD and *w*AnM also contained a suite of metabolite transport and secretion systems common to other *Wolbachia* strains that includes haem, zinc, iron (III), lipoproteins and phospholipids (Figure 3a). This conservation of pathways was also observed when the analysis was focused to only *Wolbachia* from supergroup B strains [44] (Figure 3b). In addition to these biosynthetic pathways, the Type IV and Sec-Secretion systems were also maintained in both *Wolbachia* genomes. The Type IV secretion systems (T4SS) are known to play roles in infection and survival for a diverse range of symbiotic and pathogenic intracellular bacteria [52, 53]. Both *Wolbachia* genomes contained a total of 15 T4SS related genes, organised into two operonic regions and four individual genes spread across the genome. *Wolbachia* strains in the filarial nematode *Brugia malayi* has been predicted to utilise its T4SS to secrete protein effector molecules to avoid autophagy pathways and aid in actin cytoskeleton reformation, allowing intracellular mobility [54]. Such processes may also be conserved within *Wolbachia* strains residing within insects, such as *w*AnD and *w*AnM.

**Figure 3:**
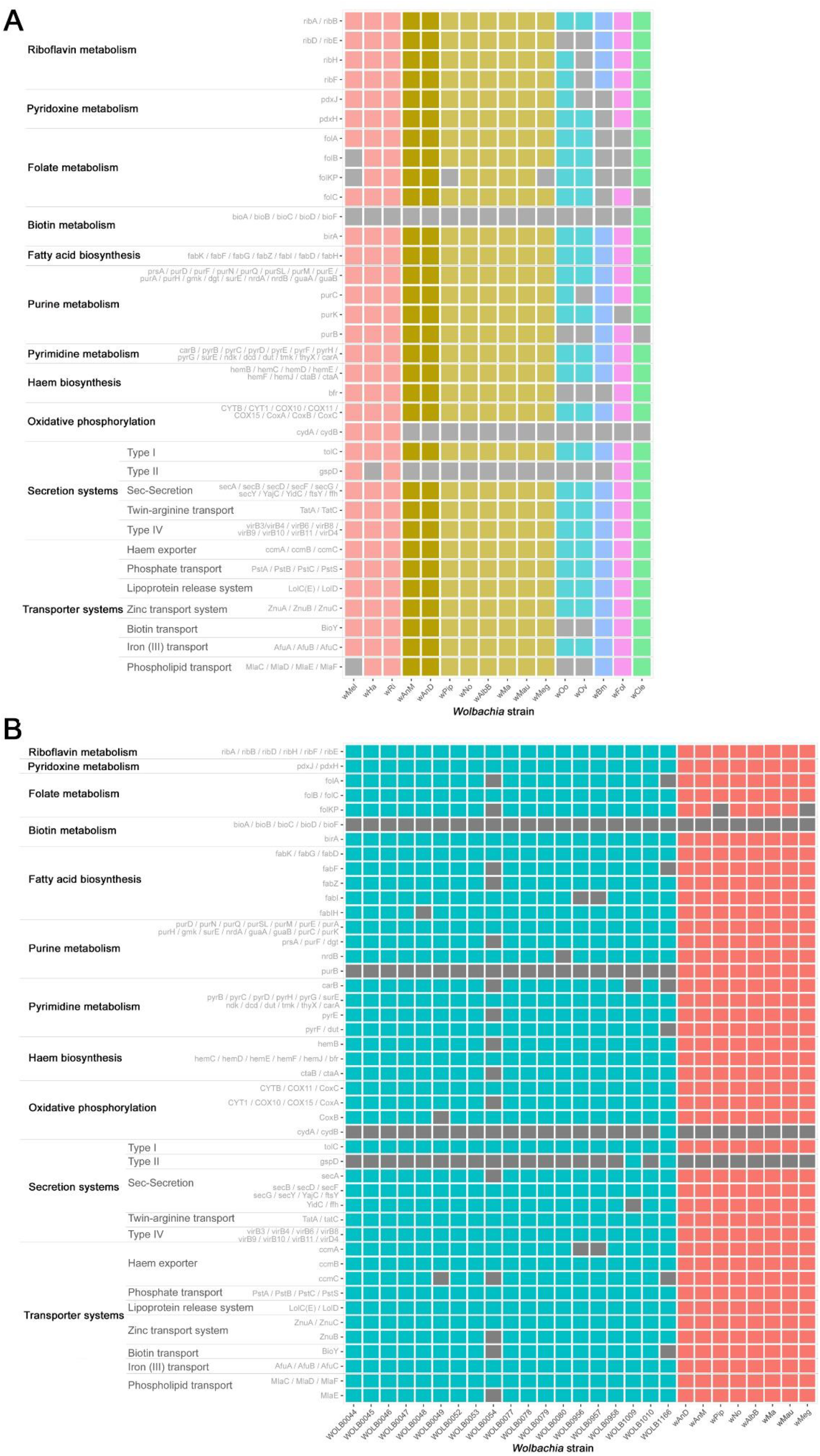
Heatmap representation of presence-absence of various genes in metabolic and secretion/transport system pathways amongst a selection of *Wolbachia* genomes. The analysed genomes are arrayed on the x-axis, with colours of the heatmap representing the various analysed supergroups. The y-axis in turn represents different genes and metabolic pathways of interest to Wolbachia studies. Greys in the heatmap represent an absence of a gene within the respective genome. **(A)** Comparative illustration of 16 near-complete *Wolbachia* genomes. Column colours are based on *Wolbachia* supergroup using a similar scheme to Figure 2, with columns representing the genomes of *w*AnD and *w*AnM are highlighted with a more intense colour of the heatmap. These genomes were observed to maintain all the genes and pathways common to supergroup B *Wolbachia*. **(B)** Comparative illustration of 29 *Wolbachia* genomes of supergroup B. Columns are coloured based on their origin, with blue columns being genomes from the study by Scholz *et al*. [44], and red being genomes from existing *Wolbachia*, including *w*AnD and *w*AnM.

### Prophage WO region identification

*Wolbachia* strains are frequently infected by a bacteriophage known as Phage WO [55], with prophage sequences predicted to be common in the genomes of *Wolbachia* strains of insects [12, 56]. These can be found in various states of completeness depending on the specific strain - some genomes are known to maintain duplicated prophage insertions that can encode a functional phage [57, 58], whilst others have been found to be degenerated [58–60]. By contrast, nematode-specific *Wolbachia* strains are known to have either no, or significantly degraded, prophage sequences [61, 62]. These prophage regions are known to maintain an EAM [63], a group of genes that encode protein domains homologous to those found in eukaryotes. This has resulted in predictions that these genes influence host-*Wolbachia* interactions by mimicking and interacting with host proteins [63]. Additionally, genes that have been implicated in the mode of action for CI have typically been found localised within these prophage EAM regions [10, 11, 15, 63].

In contrast to other *Wolbachia* strains that reside within mosquitoes, *w*AnM contained no prophage fragments identifiable via the PHASTER web server. To confirm this, we aligned the genomes of both *Wolbachia* strains from the two *Anopheles* species to their closest relative, *w*No from *D. simulans*, using the Blast Ring Image Generator (BRIG, [64]). The *Wolbachia* genome of *w*No was previously observed to have four prophage-like regions [58], ranging in size from 5.7 kbp to 47.2 kbp. Initial comparisons of the genomes showed notable gaps within the *w*AnM genome when compared to *w*No, although the same regions appear partially present in *w*AnD (Figure 4a, 4b). Overlaying coordinates for the four prophage regions that were known to be present in *w*No [58] onto this comparison, it was observed that the gaps in alignment with *w*AnM were centred on these *w*No prophage regions (Figure 4a). When the original sequencing reads were mapped to the *w*No genome, we observe very low read coverage on *w*No prophage segments (Supplementary Figure 1), whilst these reads showed even coverage of the *w*AnM genome. This indicates that these prophage regions are not present in the currently assembled genome of *w*AnM.

**Figure 4:**
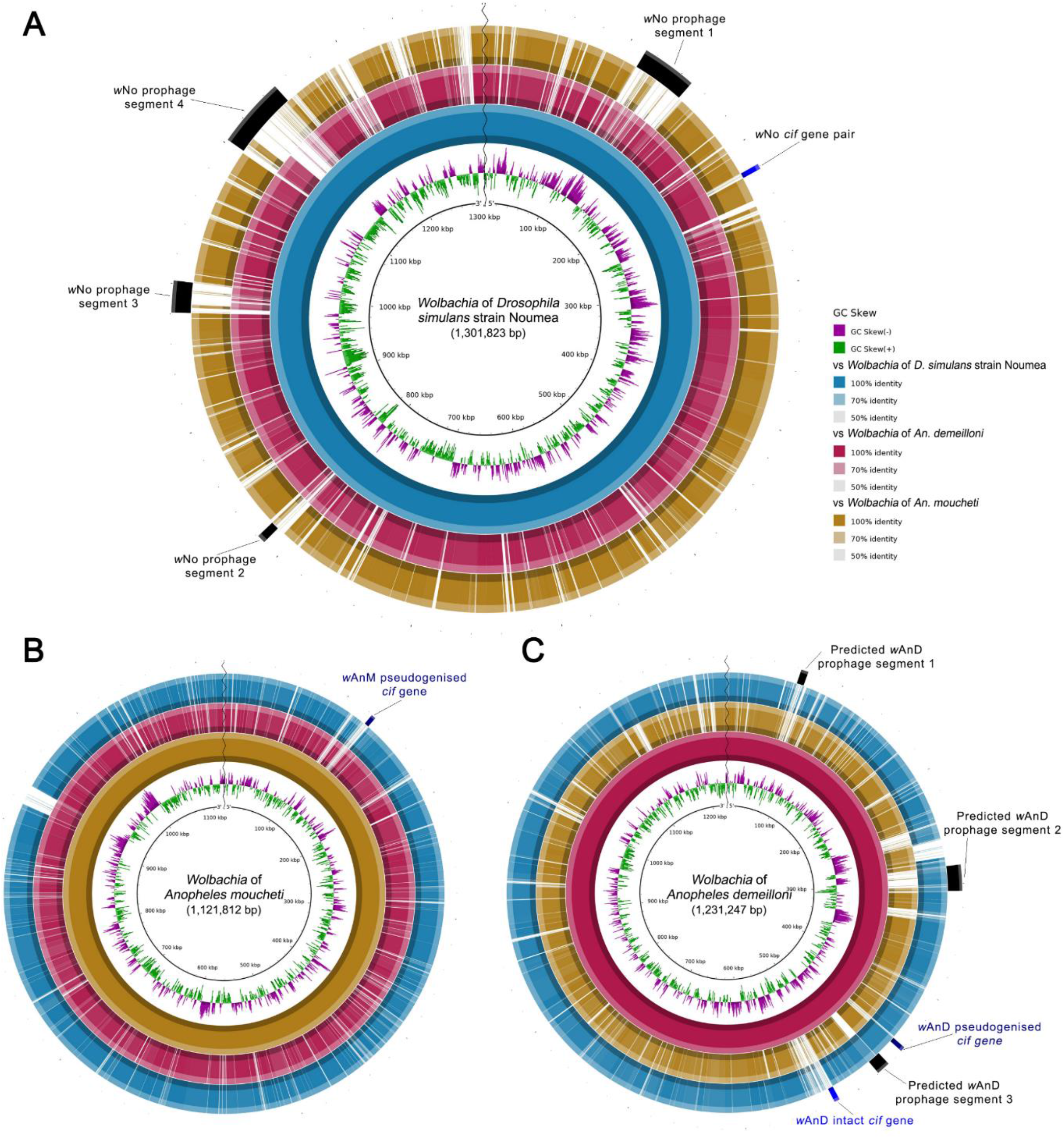
BLAST Ring Image Generator (BRIG) visualisation of prophage regions in the genomes of *w*No, *w*AnM and *w*AnD when compared to one another. Each individual ring represents the BLAST similarity score for a particular genome (represented by the different colours as defined by the key on the right of panel A) against a template (represented by the innermost solid colour ring, and the name at the centre of the circle). Differences in opacity of the rings represents the BLAST similarity score (as represented by the key on the right of panel A), with solid colours representing 100% similarity, and blank regions representing 0% similarity. The outer-most ring contains information on predicted prophage and cif gene localisations **(A)** Comparison of *w*AnM and *w*AnD against a *w*No template genome. Note how the black bars representing predicted prophage regions by Ellegaard *et al*. [58] overlap areas with no similarity against the *w*AnM genome, whilst having some similarity to the *w*AnD genome. Also note how its single, intact *cif* gene pair is located separate from previously predicted prophage regions. **(B)** Comparison against a *w*AnM template genome. Note how this genome was noted to contain no prophage regions, and its single pseudogeised cif gene pair is located in an area with no similarity to both *w*No and *w*AnD. **(C)** Comparison against a *w*AnD template genome. Predicted prophage segments 1 and 2 were predicted by the PHASTER web server, with segment 3 predicted by Blastx searches against the prophage regions WOVitA1 and WOCauB1 through to B3, as identified by Bordentstein et al. [63]. Note how of the three predicted prophage regions, two showed similarity to the *w*No genome, and one showed no similarity to either genome. Also note how its two *cif* gene pairs are located separate from, but close to, predicted prophage segment 3. Also note how the intact *cif* gene pair appears within a region that shows weak to no similarity against both *w*AnM and *w*No.

Within *w*AnD, analysis via the PHASTER web server and subsequent BlastX searches of surrounding regions identified two prophage fragments of lengths 6.3 kbp and 22.1 kbp. BlastX searches also identified an additional prophage-like region of length 11.6 kbp (Figure 4c). The total length of these prophage fragments (approx. 40 kbp) is shorter than published phage WO genomes (lengths of between 55 kbp to 65 kbp, [63, 65]). The two prophage regions identified by PHASTER is predicted to code for a total of 50 genes, 16 of which were predicted to be interrupted by either stop-codons or frameshifts. The prophage-like region identified after manual curation contained 13 genes, of which seven were predicted to be interrupted. Two of these three regions contained structural phage genes that were either intact or interrupted, with examples including phage tail, baseplate, head-tail connectors, and capsid proteins (Supplementary Figure 2).

### Cytoplasmic incompatibility (CI) factors are conserved in wAnM and wAnD

We previously reported that the genome of *w*AnD contains one intact pair of *cif* genes (JSQ73_02850, JSQ73_02855), and a second pair which showed interruptions in both genes (JSQ73_02500 through to JSQ73_02515) [36]. In turn, the genome of *w*AnM contains one pair of *cif* genes, although two internal stop codons were identified within *cifB* [36]. Phylogenetic analysis of the concatenated nucleotide sequences of *cifA* and *cifB* identified *w*AnD’s intact *cif* gene pair as clustering with the Type I group, and its pseudogenised pair clustering with the Type III group, in line with previous observations [14]. In comparison, the *cif* gene pair of *w*AnM clusters with the Type II group (IYZ83_00740 through to IYZ83_00755, Figure 5).

**Figure 5:**
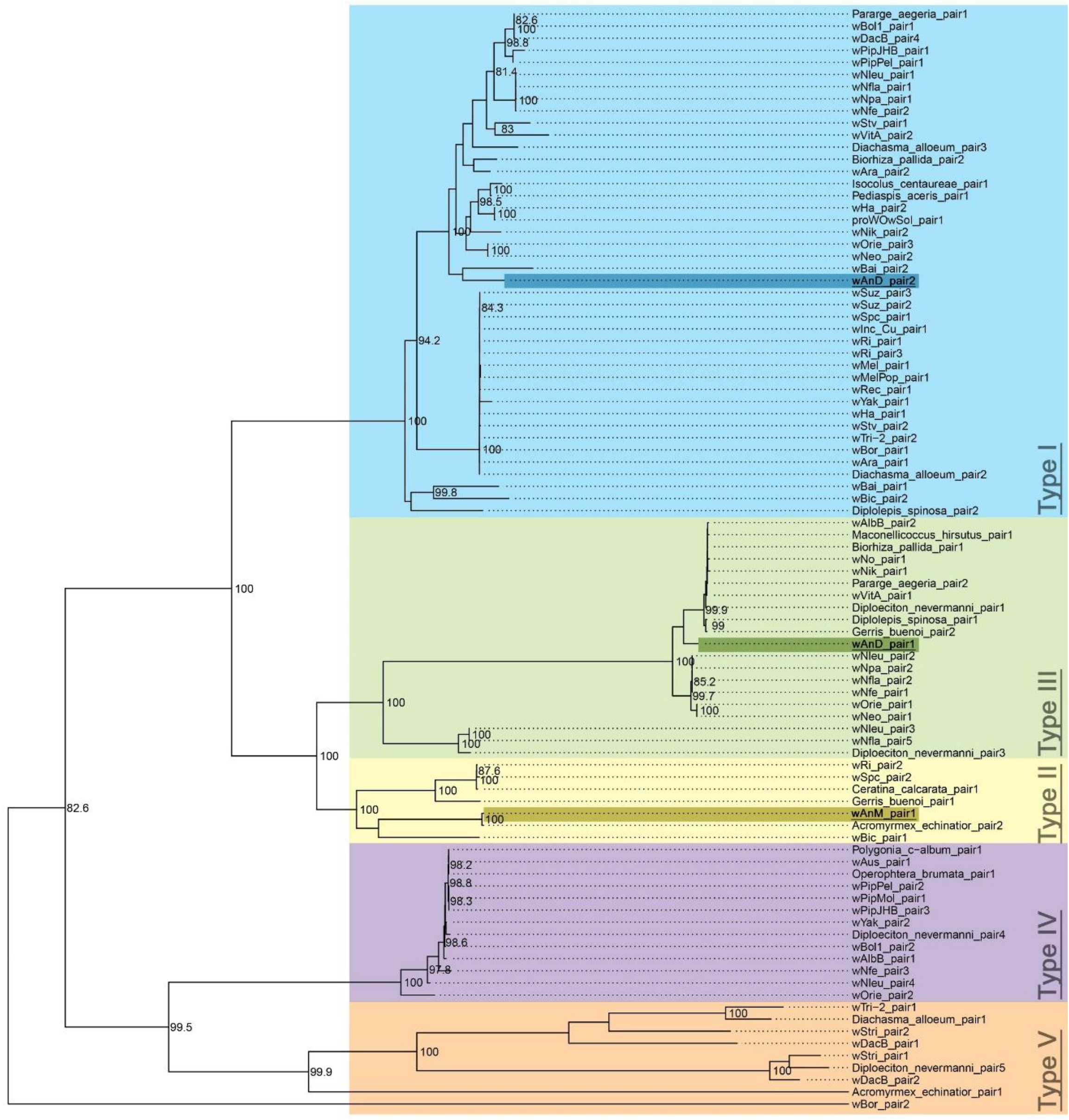
Maximum likelihood phylogenetic tree of concatenated *cif* gene nucleotide alignments, built following the methods of Martinez *et al*. [14] with 1,000 bootstrap replicates. Only bootstrap values of over 80% are shown. The five types’ of concatenated *cif* genes are highlighted with different colours, and their corresponding types annotated on them. Tree is midpoint-rooted. The two pairs of *cif* genes of wAnD were previously noted to be members of Type I and Type III, which is confirmed by this repeated analysis. The pair of *cif* genes in wAnM can be found in the well-supported Type II clade.

Within *w*AnD, the interrupted *cif* genes were of a combined 3.6 kbp in length and were located upstream of one of the prophage regions identified by the PHASTER web server (Figure 4c). Following this, the intact *cif* genes of *w*AnD combined measured 6.0 kbp in length, and is approximately 69.5 kbp downstream of the interrupted *cif* genes (Figure 4c). By contrast, the interrupted *cif* genes of wAnM were of a combined 3.6 kbp in length (Figure 4b). Interestingly, none of the three identified pairs of *cif* genes within *w*AnD and *w*AnM were located directly next to or within prophage regions, although the two within *w*AnD are located close to one (Figure 4c). This is similar to *w*No, whose single intact pair of *cif* genes were observed to be separate from predicted prophage regions (Figure 4a). Additionally, it should be noted that the *cif* gene pair of *w*AnM appear to be a unique insertion that is also not present in *w*AnD (Figure 4b).

## Discussion

This study provides a comprehensive analysis of two *Wolbachia* strains recently identified within *Anopheles* mosquitoes. Their high density and prevalence rates within field populations provides an opportunity to better understand *Wolbachia*-host interactions, as well as providing a potential tool to aid in interrupting the spread of *Plasmodium* parasites. One of the first observations from this study is that the *Anopheles*-infecting *Wolbachia* strains are not monophyletic with other *Wolbachia* strains from mosquitoes (*w*AlbB and *w*Pip). Instead, both *w*AnD and *w*AnM were located within a clade that includes several *Wolbachia* strains found within *D. simulans* and *D. mauritiana*. There have been multiple studies that show horizontal transmission of *Wolbachia* occurs regularly [47, 48], and is even possible via a plant intermediate [66]. This potential for horizontal transmission is further emphasised by a recent survey that assembled over 1,000 *Wolbachia* genomes from existing sequence data [44]. These genome assemblies are primarily distributed across various *Wolbachia* strains from supergroups A and B, whilst also generating multiple *Wolbachia* assemblies from the same host [44]. This study observed how closely related *Wolbachia* strains can be found in taxonomically unrelated hosts, as well as finding no meaningful phylogenetic clustering of different hosts and their corresponding resident *Wolbachia* strains. Such observations are similar with what is observed here with the whole-genome phylogeny of *w*AnD and *w*AnM, in relation to the wider supergroup B strains and their insect hosts.

When compared against these other sequenced *Wolbachia* strains, analysis of the *w*AnD and *w*AnM strains indicate that they maintain relatively small genome sizes for strains found within insects. Despite reduced genome sizes, both the *w*AnD and *w*AnM strains maintain similar metabolic and transport pathways found in other *Wolbachia* strains. Additionally, no biosynthetic pathways were identified that could indicate a previously unknown feature acquired in these two strains found in *Anopheles* mosquitoes. Known pathways of relevance for *Wolbachia* include haem and nucleotide biosynthetic pathways [67], as well as transport components such as the Type IV secretion system for secreting potential protein effectors [52]. The observation of smaller genome sizes could be attributed to a reduced number of mobile elements, specifically prophage regions, when compared to other *Wolbachia* strains that reside in mosquitoes such as *w*AlbB [59] and *w*Pip [57].

Following on from this, it is interesting to see how *w*AnD has degenerated prophage regions in comparison to its closest relative *w*No, whilst *w*AnM lacks prophage regions entirely. This is despite the presence of *cif* genes within both genomes, which are separate from any prophage regions, contrary to previous observations and expectations for these two features to be co-localised [10, 11, 24, 63, 68]. However, this separation of *cif* genes is not unique to just these two *Wolbachia* strains in *Anopheles*, but also to the closely related *Wolbachia* strains *w*No, *w*Ma, and *w*Mau (the first infecting *D. simulans*, the latter two *D. mauritiana*), which have been shown to maintain *cif* genes that are distinctly separate from any prophage WO region [45, 69] (Supplementary Figure 3). It is tempting to speculate that this separation of *cif* genes and prophage regions be a unique feature of this clade of *Wolbachia*. For comparison, the genomes of both *w*AlbB and *w*Pip maintain *cif* genes that are associated with prophage WO regions [10, 59]. In addition to this separation from prophage regions, both strains *w*Ma and *w*Mau were observed to have an interrupted *cif*B gene [45, 69], similar to what is observed in *w*AnM, and are both incapable of inducing CI, but capable of rescuing it, when crossed with *w*No-infected mates [70, 71]. Unlike *w*AnM however, all three of *w*No, *w*Ma and *w*Mau‘s *cif* gene pairs are found within the Type III phylogenetic group, whereas the *cif* gene pair identified in *w*AnM can be placed within Type II, which is unique amongst this group of five *Wolbachia*. BRIG comparisons of the different genomes appear to indicate this *cif* gene pair to be a unique insertion. Furthermore, whilst *w*AnD’s degenerated *cif* gene pair was noted to be a member of the Type III group, its intact *cif* gene pair also appears unique among this group of *Wolbachia* as a member of Type I. Like *w*AnM’s sole *cif* gene pair, this intact *cif* gene appears to be a unique insertion event, separate from prophage elements.

How such insertion events within both *w*AnD and *w*AnM have come to happen, and where they have come from, is currently an open question that warrants further investigation, alongside how this group of *Wolbachia* maintain *cif* gene pairs that appear separate from identifiable prophage WO regions. One possible explanation is that the recent ancestors for these strains of *Wolbachia* may have acquired these *cif* genes from a recent phage WO insertion that has very recently become degenerated [69]. Alternatively, these prophage regions could have been removed from the genome by phage excision events. Previous publications have discussed what could happen to the *cif* gene pairs, as well as the *Wolbachia* which carry them, once CI is no longer able to induce evolutionary pressure on their hosts [72–75]. For instance, a recent survey of CI genes in *Wolbachia* predicted how, without evolutionary pressure, these CI genes would likely degrade over time, starting with *cif*B, the ‘toxin’ component of the phenotype, followed by *cif*A, the ‘antidote’ component [14]. Alternatively, it has also been suggested that the degradation of the *cif* genes may be related to the absence of prophage regions [14, 15], with the former being an adaptation used by the latter to spread within *Wolbachia* populations. Thus, once the prophage regions are removed, it is predicted that the *cif* genes, and thus the CI phenotype, will have no evolutionary pressure to maintain themselves within *Wolbachia* [14, 15]. We observe this occurring to some degree in this study, with the dissociation of prophage regions from the *cif* genes, the interrupted Type III pair observed in *w*AnD, and how *w*AnM carries interruptions in its Type II *cif*B gene specifically. Once the phenotype these *Wolbachia* strains exert on their hosts can be properly elucidated, a longitudinal study on the *cif* genes within them is imperative. The results of such study could allow for further insights into *Wolbachia* biology and the evolution of the CI phenotype.

Despite the questions as to how this may have occurred, the observed similarities and differences between *w*AnM and its related strains *w*Ma, *w*Mau and *w*No are intriguing, and raises the possibility that *w*AnM may not cause CI in its *Anopheles* host. This is perhaps counterintuitive, considering the high, but variable, prevalence rates of *w*AnM in field populations of *A. moucheti* [36, 42]. This prevalence rate is a feature shared with *w*AnD [36, 42], which is more likely to be capable of inducing CI due to the presence of intact *cif* genes from the Type I group, of which *w*Mel shares. For comparison, our previous work had shown the prevalence rates of *w*AnM to be between 17.5% and 75%, which is slightly lower than *w*AnD prevalence rates, shown to be between 38.7% and 100%. Yet the ability for *Wolbachia* to persist in ppulations without inducing CI is known, as there are instances of *Wolbachia* which stably infect host populations without any overt reproductive parasitism phenotype [8, 76, 77]. Explanations for this have focused on *Wolbachia* providing some form of fitness benefits to their host. For instance, the Wolbachia strain *w*Au of supergroup A spread through lab-based, uninfected host populations of *D. simulans* without inducing CI [78]. This persistence of *w*Au could be linked to an ability to induce protection against viral infections [79], and it is tempting to speculate that *Wolbachia* may provide protection against pathogens of the mosquito. While such studies focus on *Wolbachia* of supergroup A, there has been some evidence that *w*Mau of supergroup B may also confer a fitness benefit for their host via stimulating egg production [69, 80]. Further research will still need to be done to confirm if *w*AnM can confer similar fitness benefits to its host, or have the potential to inhibit *Plasmodium* or viruses, and whether host-*Wolbachia* backgrounds may play any role in this.

The identification of natural *Wolbachia* infections in *Anopheles* shows promise for future control strategies of *Plasmodium* parasites. Whilst these strains show no pathways that are uniquely present or absent, they do exhibit unusual genomic arrangements with regards to the presence of prophage and *cif* genes. This has potential implications on their relationship with their respective Anopheline hosts, potentially making them good candidates for transinfection into other medically relevant *Anopheles* species, such as *A. gambiae* s.s. Further studies would be required to fully examine these *Wolbachia* strains and elucidate their predicted phenotypes of CI and pathogen blocking, both in the context of natural and artificial associations.

## Methods

### Sequence data collection and genome quality assessment

Both genomes assemblies of *w*AnD and *w*AnM were manually curated (i.e. gaps, indels and synteny) using the approach described by Tsai and colaborators 2010 [81], Mummer/Nucmer software tool v4.0.0 [82], Mauve v2.4.0 [83] and Tablet v1.21.02.08 [84]. To complement the genomes of *w*AnD and *w*AnM [36], whole genome sequences of 15 *Wolbachia* genomes were downloaded from the National Centre for Biotechnology Information, with these genomes spanning supergroups A through to F (full information available in supplementary table). An additional 25 *Wolbachia* genomes were also downloaded from the European Nucleotide Archive (ENA). These additional genomes were sequenced as part of a large-scale study [44] which looked at assembling *Wolbachia* genomes from a variety of existing sequencing data of various insects. For the 16 published *Wolbachia* genomes downloaded from NCBI, Refseq annotations were obtained via their PGAP [49]. The 25 genomes downloaded from ENA were also annotated using a local installation of PGAP [49], with an additional seven genomes of *Wolbachia* from supergroup B downloaded from NCBI also annotated using this local installation for consistency [49]. The genomes of both *w*AnD and *w*AnM were annotated using both methods. All genome accession numbers used in this study, as well as a summary of their annotations used in this study are provided in supplementary table.

To confirm genome completeness, nucleotide sequences of all downloaded genomes, as well as the assembled genomes of *w*AnD and *w*AnM, were used as input into the programs BUSCO (v5.0.0, [43]) with the lineage option set to “rickettsiales_odb10”. This program analyses genome completeness via comparison against a selection of marker genes (total 364 genes) predicted to be present in single copies based on the input genome’s lineage. Genomes that showed significantly lower completeness levels, (less than 80% completeness) were excluded from orthologue and pathway analyses. This resulted in six of the *Wolbachia* genomes [44] to be removed from these additional analyses.

### Phylogenetic, pangenome and metabolic pathway analysis

A total of 34 *Wolbachia* genomes were used for phylogenetic analysis of supergroup B *Wolbachia* (genomes of *w*AnD and *w*AnM, seven *Wolbachia* genomes from NCBI, and 25 from ENA). These genomes were used as input into the program wgsim ([86, 87], version 1.9), which simulates sequencing reads from a genomic template. Base error, mutation, fraction of indels and indel extension probability were set to zero, read lengths set to 100 and a total of ten million reads simulated for each genome. These simulated reads were then used to generate a Single-Nucleotide Variant (SNV) alignment via Snippy v4.6.0 [88] using the *w*No genome as reference (genome accession number GCA_000376585.1). Gubbins v3.0.0 [89] was used for removing recombinant events. Recombination-free alignment of all 34 genomes was then analysed with IQTree v1.6.12 [90] using default parameters, with a GTR substitution model using 1000 non-parametric bootstrap replicates for branch support.

### Orthologous group detection

Orthologous group detection was performed in two separate parts-first was to compare coding protein sequences amongst *Wolbachia* of supergroup A through to F, whilst the second was to compare coding protein sequences amongst *Wolbachia* of supergroup B specifically.

For orthologue analysis amongst *Wolbachia* of supergroup A through to F, Refseq protein annotations for the 15 genomes downloaded from NCBI were used, alongside Refseq protein annotations for *w*AnD and *w*AnM. This list of 17 protein sequences were used as input into the program OrthoFinder ([91], v2.5.1) using default parameters. Orthogroups that were common or unique between all 17 *Wolbachia* strains were subsequently plotted using the R program package ‘UpsetR’ ([92], v1.4.0). Additional querying of the data was then performed using the R program package ‘ComplexHeatmaps’ ([93], v2.5.5).

For orthologous group detection amongst a wider selection of supergroup B *Wolbachia*, a total of 31 *Wolbachia* genomes were used (seven from NCBI, 22 from ENA), alongside the assembled genomes of *w*AnD and *w*AnM (supplementary table). Protein gene annotations for all *Wolbachia* genomes from a local installation of the NCBI PGAP ([49], build5508 2021-07-01) were used as input into the program Orthofinder ([91], v2.5.1) using default parameters. Orthogroups were again visualised using the R program UpsetR ([92], v1.4.0), with additional data querying performed using the R program ComplexHeatmaps ([93], v2.5.5). Genes of interest identified within these orthogroups, e.g. those that were unique to particular genomes, were further analysed using the PFam website’s sequence search [94, 95] and NCBI’s BlastP [96]. Comparison of the identified nucleotide regions that had similarity to the *Osmia lignaria* gene XP_034172187.1 was performed by BlastN and BlastX, with visualisations performed using Easyfig [97].

### Construction of metabolic pathways

The genomes of both *w*AnD and *w*AnM were submitted to the NCBI Prokaryotic Annotation Pipeline, with a GenBank Flatfile being generated as a result. This flatfile was then downloaded, and used as input into BioCyc’s Pathway Tools program ([98], v24.0) and Pathologic ([99, 100], v24.0). Pathologic is able to assign protein function and pathways to annotated genes based on name and/or automated blast hits. To address proteins with ‘ambiguous’ function within metabolic pathways, all predicted protein-coding genes of both *w*AnD and *w*AnM were submitted to the EggNOG online server, which allows for the automated transfer of functional annotations ([101], v2.0.1). Predicted protein-coding genes were also submitted to the KEGG Automatic Annotation Server (KAAS, [51], last updated April 3rd 2015) as a second method for functional annotation. Any proteins identified by Pathologic as having an ‘ambiguous’ function was then manually cross-checked with the outputs of EggNOG and KAAS, and enzyme code numbers assigned. This process was repeated for a selection of *Wolbachia* genomes from supergroup B (supplementary table). Once this process was completed, Pathway Tools’ Pathway Overview and Comparison options were then used to compare pathways between the different *Wolbachia* strains. A selection of these biosynthetic and transport pathways was then made, based on prior literature investigating their importance to the *Wolbachia*-host endosymbiotic relationship. Gene presence and absence within these pathways was then manually scored, and plotted out into a heatmap using R’s GGplot2 package [102].

### Characterisation of Cytoplasmic Incompatibility Factor genes

Phylogenetic placement of the three sets of *cif* gene pairs from both *w*AnD and *w*AnM were made following the methods of Martinez *et al.* [14]. Briefly, nucleotide sequences for Cytoplasmic Incompatibility Factor (*cif*) A and B genes for all five monophyletic types were obtained from the supplementary materials of Martinez *et al.* [14]. Partially sequenced *cif* genes were discarded, and the nucleotide sequences for *cif*A and *cif*B genes were aligned separately using the program MAFFT. The separate alignments were then used as input into the online GBlocks server ([103], v0.91b) with default ‘stringent’ parameters to filter out weakly conserved regions of the alignment. Once filtering was done, the separate nucleotide sequence alignments were then concatenated using Seqkit ([104], v0.15.0), and used as input for PhyML ([105], v3.0), using the GTR GAMMA substitution model of evolution and 1,000 bootstrap replicates. The outputted newick formatted tree was then annotated using the GGTree package in R ([106], v2.2.4).

### Ankyrin, Prophage and IS element detection

Ankyrin domains were detected using five HMMer profiles ([95], ID numbers PF00023.31, PF12796.8, PF13606.7, PF13637.7, PF13857.7). These profiles were generated via first downloading associated alignment files from the PFAM protein database [94] as Stockholm formatted seed files. The HMMer suite ([95], v3.1b2) was then used to build HMM profiles from these seed files. These profiles were then compared against the protein amino acid sequences annotated from *w*AnD and *w*AnM to identify any protein-coding genes containing an ankyrin domain. This analysis was then repeated for a selection of *Wolbachia* genomes to allow for direct comparisons to be made.

Prophage sequences were identified within the genomes of *w*AnD and *w*AnM using the PHASTER web server [107]. Assembled contig sequences of both genomes were uploaded separately to the server, checking the option to note that the input consists of multiple separate contigs. In the case of *w*AnD where prophage regions were detected, results were downloaded and manual curation of the identified prophage regions was performed using the Artemis genome browser [108] to identify prophage genes overlapping these regions. Additional BlastX searches were performed on neighbouring genes against Phage WOVitA1 sequences (GenBank genome reference HQ906662.1) to screen for genes that may be associated with prophage WO’s eukaryotic association module.

Insertion sequence element detection was performed by separately concatenating the contigs of *w*AnD and *w*AnM, resulting in two contiguous genomes. These were then submitted to ISSaga ([109], v1.0) and results tables obtained. Manual curation was then performed using the original contigs for both *Wolbachia* genomes, with true-positive IS elements called by comparison of annotations from ISSaga, PGAP annotation, and BLAST searches against the ISFinder database.

## Author statements

S.Q. designed methodology, conduct investigation and formal analysis, designed visuals, and wrote the original draft. L.C. designed methodology, conduct investigation and formal analysis, and wrote the original draft. C.L.J. conceptualised the study, conducted investigation and resource collection, and wrote the original draft. S.T. designed methodology and software. T.W. conceptualised the study, conducted investigation and resource collection, secured funding, and wrote the original draft. G.L.H. conceptualised and supervised the study, secured funding, and wrote the original draft. E.H. conceptualised and supervised the study, designed the methodology, and wrote the original draft. All authors have read and approved the final manuscript.

The authors declare that there are no conflicts of interest.

## Funding information

EH and GLH acknowledge support from BBSRC grant BB/V011278/1. ST was supported by the Wellcome SEED award 217303/Z/19/Z to EH. LC was supported by NIAID R01-AI116811. TW and CLJ were supported by a Sir Henry Dale Wellcome Trust/Royal Society fellowship awarded to TW (<u>101285</u>): https://wellcome.org and https://royalsociety.org. TW was also supported by a Royal Society challenge grant (CHG\R1\170036). GLH was also supported by the BBSRC (BB/T001240/1), a Royal Society Wolfson Fellowship (RSWF\R1\180013), the NIH (R21AI138074), the EPSRC (EP/V043811/1), the UKRI (20197 and 85336), and the NIHR (NIHR2000907).

## Notes

https://figshare.com/projects/Wolbachia_endosymbionts_in_two_Anopheles_species_indicates_independent_acquisitions_and_lack_of_prophage_elements/126533

### Competing Interest Statement

The authors have declared no competing interest.

