## Supplementary figures and images for "*Wolbachia* endosymbionts in two *Anopheles* species indicates independent acquisitions and lack of prophage elements"

### Supplementary figure 1

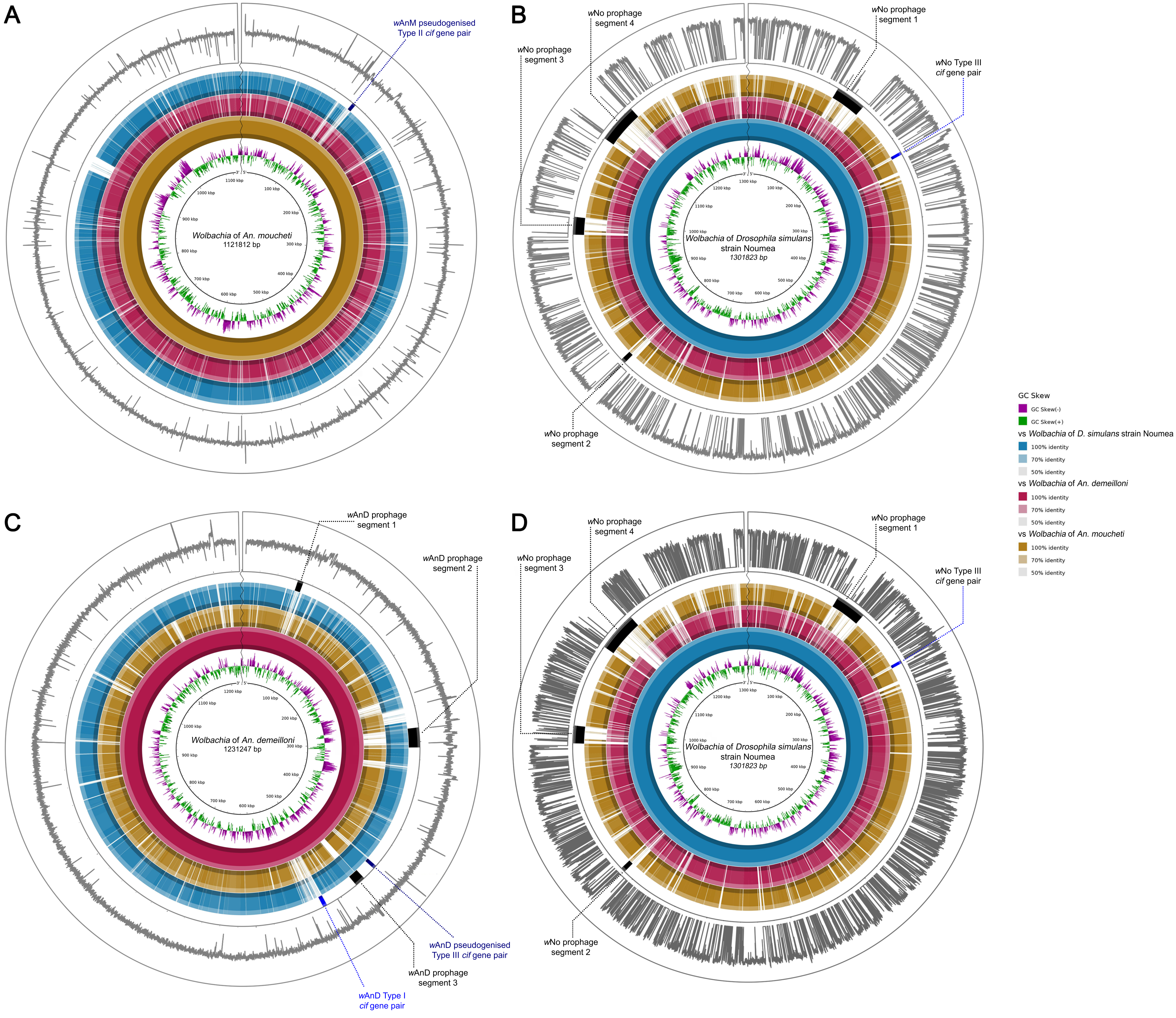

### Supplementary figure 2

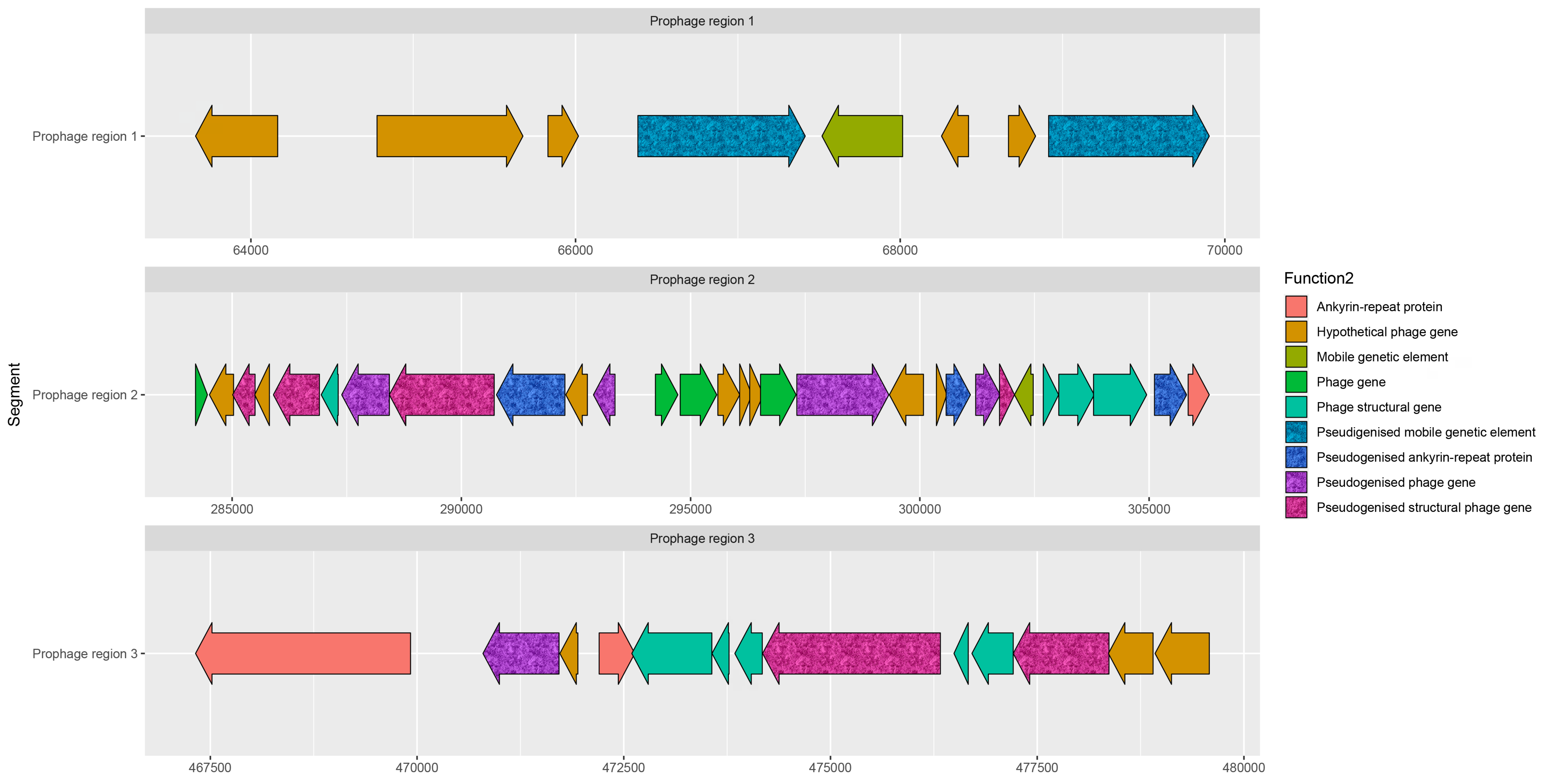

### Supplementary figure 3

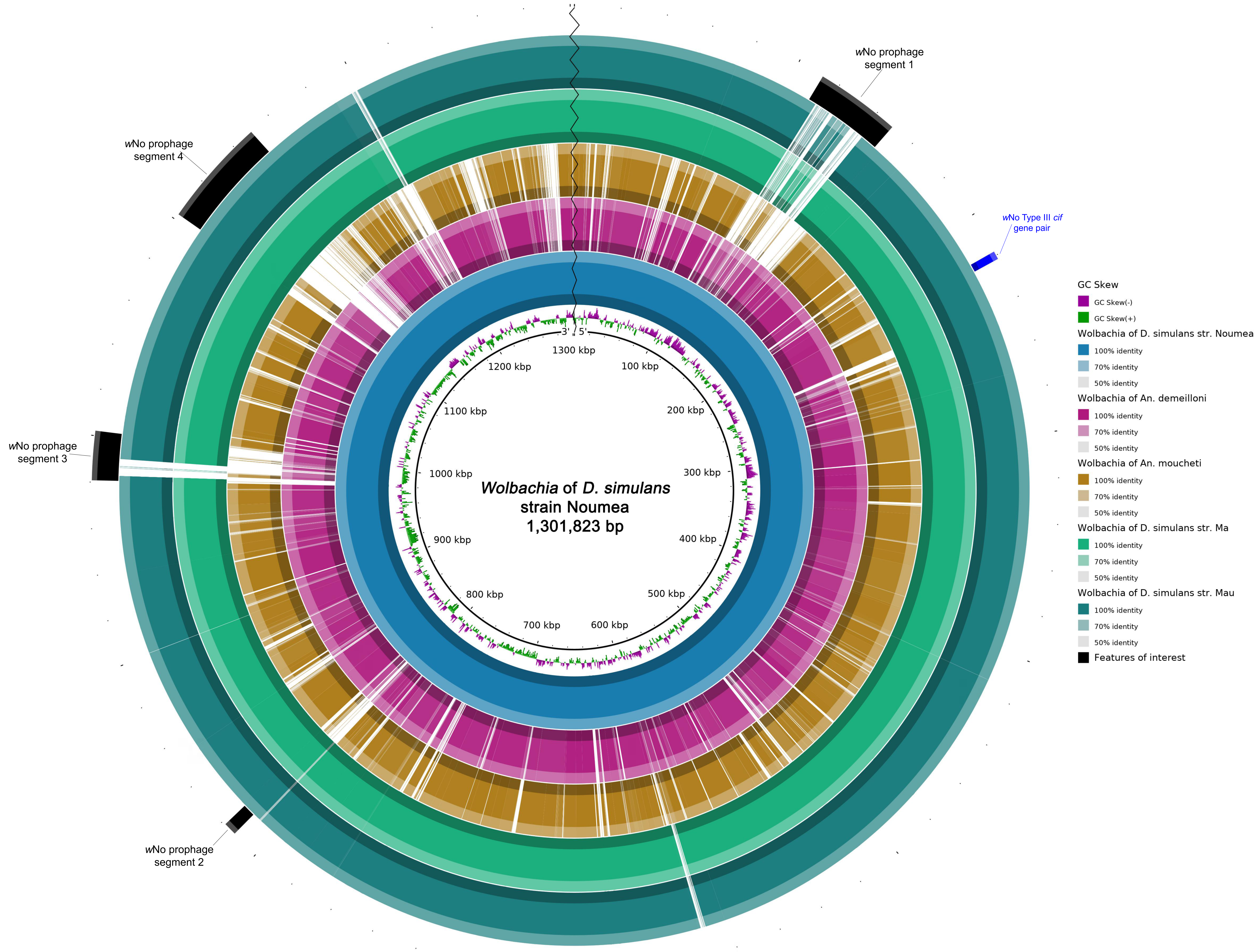
